# Ant guards influence the mating system of their plant hosts by altering pollinator behaviour

**DOI:** 10.1101/2020.02.11.943431

**Authors:** Nora Villamil, Karina Boege, Graham N. Stone

**Affiliations:** Institute of Evolutionary Biology, University of Edinburgh, Ashworth Laboratories, The King’s Buildings, Edinburgh, EH9 3FL, United Kingdom; Instituto de Ecología, Universidad Nacional Autónoma de México. A.P. 70-275. Ciudad Universitaria, C.P. 04510, Ciudad De México. México; Department of Ecology and Evolution, Université de Lausanne, Biophore, CH-1015, Lausanne, Switzerland

**Keywords:** male fitness, mating systems, myrmecophily, non-pollinators, outcrossing, plants

## Abstract

Ant guards can increase plant fitness by deterring herbivores but may also reduce it by interfering with pollination, hence ant-plant interactions are ideal systems in which to study costs and benefits of mutualisms. While ant impacts on herbivory are well-studied, much less is known about impacts on pollinators and associated consequences for plant mating systems and fitness. We used field experiments to quantify the effect of ant guards on pollinator community composition, frequency and duration of flower visits, and cascading effects on plant mating system and plant fitness in *Turnera velutina* (Passifloraceae). Although ant patrolling did not affect pollinator community composition or visitation frequency, it decreased pollinator foraging time and flower visit duration. Such behavioural changes resulted in reduced pollen deposition on stigmas, decreasing male fitness whilst increasing outcrossing rates. This study contributes to understanding how non-pollinators, such as these defensive mutualists, can shape plant mating systems.

## Introduction

Ant-plants are excellent systems in which to explore the costs and benefits of multispecies mutualisms. While aggressive ants increase plant fitness by defending their host plants from herbivores (Bentley 1977; Beattie 1985; Martin & Doyle 2003), they can also decrease plant fitness disrupting plant–pollinator mutualisms and repelling other plant-beneficial predatory arthropods (Koptur *et al.* 2015). Ants can disrupt pollination by consuming floral structures, damaging pollen (Stanton *et al.* 1999; Frederickson 2009; Stanton & Palmer 2011; Dutton & Frederickson 2012; Malé *et al.* 2015), or deterring flower visitation by pollinators (Assunção *et al.* 2014; Villamil *et al.* 2018). Yet, information on their impacts on pollinators, pollen transfer and seed set is still limited, and only few studies have addressed the ecological costs of ants via pollinator deterrence (Romero & Koricheva 2011).

Ant aggressivity may be a double-edged sword underlying the core ecological costs and benefits of myrmecophily. More aggressive ants may be better defenders against herbivores, but may also pose a higher predation risk to other mutualistic guilds such non-ant predators of herbivores or pollinators (Ness 2006; Ohm & Miller 2014; Jones & Koptur 2015; Villamil *et al.* 2018). Furthermore, the metrics commonly used in quantifying the effectiveness of indirect defences against herbivores – reduction in damage by herbivores – are unlikely to reveal pollination-associated impacts on plant fitness (Dukas 2001; Gaume *et al.* 2005; Ness 2006; Goncalves-Souza *et al.* 2008; Frederickson 2009; Romero & Koricheva 2011; Stanton & Palmer 2011; Dutton & Frederickson 2012; Ohm & Miller 2014; Jones & Koptur 2015; Malé *et al.* 2015) and a multispecies approach is needed to improve our estimates of the net outcomes of mutualistic interactions.

Ant impacts on pollinators may be consumptive (through predation) or non-consumptive, defined as changes in prey traits or behaviours in response to perceived predation risk (Preisser *et al.* 2005; Sheriff & Thaler 2014). The magnitude of non-consumptive effects on pollinator and plant fitness can be similar to, or higher than, that of direct consumptive effects (Preisser *et al.* 2005; Romero *et al.* 2011; Clinchy *et al.* 2013; Sheriff & Thaler 2014). However, the mechanism(s) by which predators influence pollinator behaviour and impact on plant fitness remain entirely unknown for the majority of ant-plants (Romero & Koricheva 2011).

The few studies on the effects of ant patrolling on pollinator behaviour suggest that ants can have positive or negative consequences for plant fitness. For example, lower seed set in *Ferocactus wislizenii* plants tended by aggressive ants was attributed to a three-fold reduction in pollinator visitation frequency (Ness 2006). However, this hypothesis was not experimentally tested. Alternatively, an increase in fruit set in ant-patrolled plants of *Psycothria limonensis* was attributed to pollinator relocation, where ant threats might have caused pollinators to spend less time per flower and visit more flowers, promoting pollen transfer (Altshuler 1999). Again, this mechanism was inferred, but not experimentally tested. Previous experiments on *Turnera velutina* showed that ant corpses placed inside flowers reduce pollinator visit duration (Villamil *et al.* 2018). However, such an experimental setup may differ from natural circumstances as flower occupation by ants is a rare event, and live ants (in contrast to dead ones) do not remain immobile in the flowers for long periods. Overall, the presence of ants can promote changes in pollinator community composition, visit frequency and duration. This, in turn, could drive positive or negative impacts of ant-pollinator interactions on plant reproduction (Altshuler 1999; Ness 2006). To date, no study has quantified the impacts of ant patrolling on pollinator visitation behaviour, plant mating systems, and fitness under natural conditions.

We estimated the ecological and potential evolutionary consequences of myrmecophily on the pollination biology, mating system, and fitness of *Turnera velutina* (Passifloraceae), a self-compatible ant-plant using an ant exclusion field experiment. We addressed the following questions: (i) What is the effect of ant patrolling on pollinator visitation? (pollinator community composition, visitation frequency, duration, and behaviour) (ii) Does ant patrolling affect the host plant mating system? (iii) Does ant patrolling affect pollen transfer dynamics? (iv) Does ant patrolling affect plant male fitness? First, because smaller or solitary pollinator taxa are expected to be more vulnerable to predation risk than larger or social species (Dukas & Morse 2003) (Clark & Dukas 1994; Abbott & Dukas 2009) we predicted that the pollinator community composition on ant-excluded plants should be biased, relative to plants with ants, towards smaller and solitary taxa. Second, due to ant-associated predation risk, we hypothesised that flowers of ant-occupied plants would receive fewer and shorter pollinator visits, with higher rates of flower avoidance (pollinator failure to land). Last, we had three different predictions for the effects of ants on pollinator visitation and plant mating system, depending on the magnitude of ant-related impacts on pollinator behaviour: (a) Ants strongly deter pollinators, leading to reduced visitation frequency, shorter visits, pollinator limitation, and reduced seed set in ant-occupied plants. (b) Ants partially deter pollinators forcing them to relocate to other flowers within the same plant, leading to higher visitation frequency but reduced visit duration and higher rates of geitonogamy (intra-plant pollination) in ant-occupied plants. (c) Ants partially deter pollinators, forcing them to relocate to flowers of different plants, leading to higher visitation frequency but reduced visit duration, and higher outcrossing rates (inter-plant pollination) in ant-occupied plants, increasing seed genetic diversity.

## Materials and methods

### Study site and system

Field experiments were conducted in coastal sand scrub at Troncones, Guerrero, on the southern Pacific coastline of Mexico (17°47’ N, 101° 44’ W, elevation < 50 m). *Turnera velutina* (Passifloraceae) is a Mexican endemic shrub (Cuautle & Rico-Gray 2003; Arbo 2005) that establishes a facultative mutuaalism with 10 ants species in Troncones (Zedillo-Avelleyra 2017) rewarding them with extrafloral nectar (Villamil *et al.* 2013). *Turnera velutina* is a self-compatible, herkogamous species that requires pollinators for seed production (Sosenski *et al.* 2016). Although it flowers year-round, flowering peaks during summer (Cuautle *et al.* 2005) and the entomophilous flowers last one day (Sosenski *et al.* 2016). Pollinator rewards are pollen and floral nectar (Sosenski *et al.* 2016; Villamil *et al.* 2018). At Troncones, native butterflies are the dominant flower visitors of *T. velutina*, followed by the introduced honeybee (*Apis mellifera*); native bees, wasps, and occasionally flies also visit the flowers.

### Ant exclusion and experimental setup

We identified six replicate arrays of plants, each of which was at least 10 m from any other array, and comprised two large focal plants producing > 6 flowers per day, and separated by >2 m. One focal plant was randomly designated as control, with natural levels of ant-guards. The second was designated ant-excluded, excluding ants from all stems using Tanglefoot™ (Fig 1a). Both focal plants in each array were isolated from other plants by trimming or tying back any surrounding vegetation. Exclusion treatments were checked daily and Tanglefoot™ was replenished if required. Each focal plant pair was surrounded by 6-10 neighbouring adult plants of *T. velutina* >2 m away (Fig. 1a).

**Figure 1.**
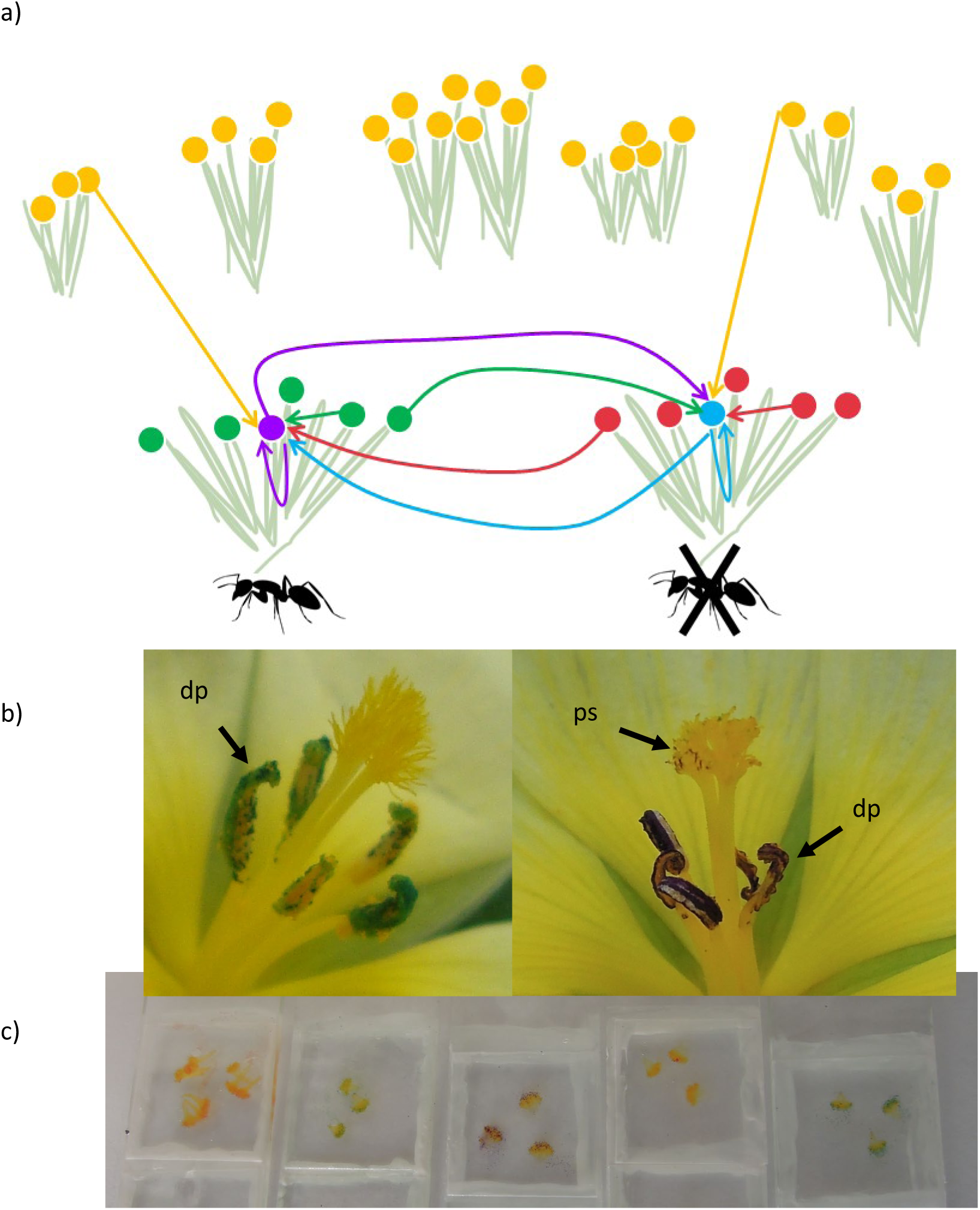
Experimental design and methods. (a) Diagram of the experimental setup showing a plant pair with or without ant patrolling. Within each experimental plant, the focal flower is represented by a uniquely coloured circle, and satellite flowers are the five circles coloured differently. The surrounding bushes with yellow circles represent undyed (naturally yellow) flowers from neighbouring plants bagged to secure allogamous pollen. (b) Photograph of dyed pollen on the anthers (dp) and dyed pollen grain on the flower stigmas (ps). (c) Photograph of stigma squash slides with dyed pollen grains.

To assess the effect of ant patrolling on pollen transfer and its consequences on the rate of selfing, geitonogamy, and outcrossing, we dyed the anthers and pollen of control and ant-excluded plants, using four contrasting dyes (red, blue, green or purple) (Fig. 1b). Within each focal plant, one flower was designated as a focal flower, whilst the other five flowers were designated satellite flowers (Fig. 1a). The anthers of the focal flower were dyed using one colour, whilst the anthers of all satellite flowers were dyed in a second colour. The remaining two colours were used on the other focal plant within the array, differentially dyeing the anthers of focal and satellite flowers (Fig. 1a). Pollen from the neighbouring non-focal *T. velutina* plants within the array was left undyed (naturally yellow-orange). The dyeing treatment was repeated in each of the six plant arrays.

Every morning before anthesis, six flower buds per plant (1 focal + 5 satellite buds) were bagged to exclude visitors. All additional pre-anthesis buds were removed to standardise floral display across focal plants. Once the corollas were fully open, anthers were dyed and flowers were re-bagged until the dye dried and anthers dehisced, exposing the dyed pollen (Fig. 1b). To ensure a minimum common supply of allogamous pollen across all flower pairs, 10-12 flowers from the neighbouring plants within the array were also bagged before anthesis and remained bagged until the visitation observations started. Stigmas from focal flowers were collected at the end of the anthesis period to count pollen grains received, as detailed below (Fig. 1c).

### Pollen dyes

Anthers of focal and satellite flowers were dyed once the corolla opened completely (∼0800-0815), but before anther dehiscence. Anthers were individually embedded in a droplet of dye until soaked, and flowers were bagged again until anthers dehisced and the released pollen was dry. The dyes used were methyl violet (purple), Green S (green), safranin (red), and methylene blue (blue) (for further details see Supplementary material). Previous studies showed that dyeing *Turnera velutina* anthers in these colours effectively dyed pollen grains had no effect on pollinator visitation (Ochoa Sánchez 2016). Towards the end of anthesis (11:30), pistils from focal flowers were collected in Eppendorf tubes and slide mounted as a glycerine squash (Kearns & Inouye 1993; Ochoa Sánchez 2016).

### Pollinator visitation

We recorded pollinator visitation to all six flowers on control and ant-excluded focal plants. Every focal plant was observed for two 20-minute periods – one immediately after bag removal when flowers had a full pollen and nectar load and the second 90 minutes later. Flowers remained bagged until their first observation round started to ensure all flowers had a full pollen and nectar load. We recorded the identity, frequency, duration, and behaviour of floral visitors and visits, as detailed below. We conducted a total of 40 h of observations of 360 flowers on 12 plants over five days. Statistical analyses were conducted in R version 3.5 (R Core Team 2016). All mixed effects models were fitted using ‘lme4’ R package (Bates *et al.* 2016) and *post-hoc* Tukey comparisons were tested using the ‘multcomp’ R package (Hothorn *et al.* 2008). All model specifications are reported in detail in Table 1.

**Table 1.**
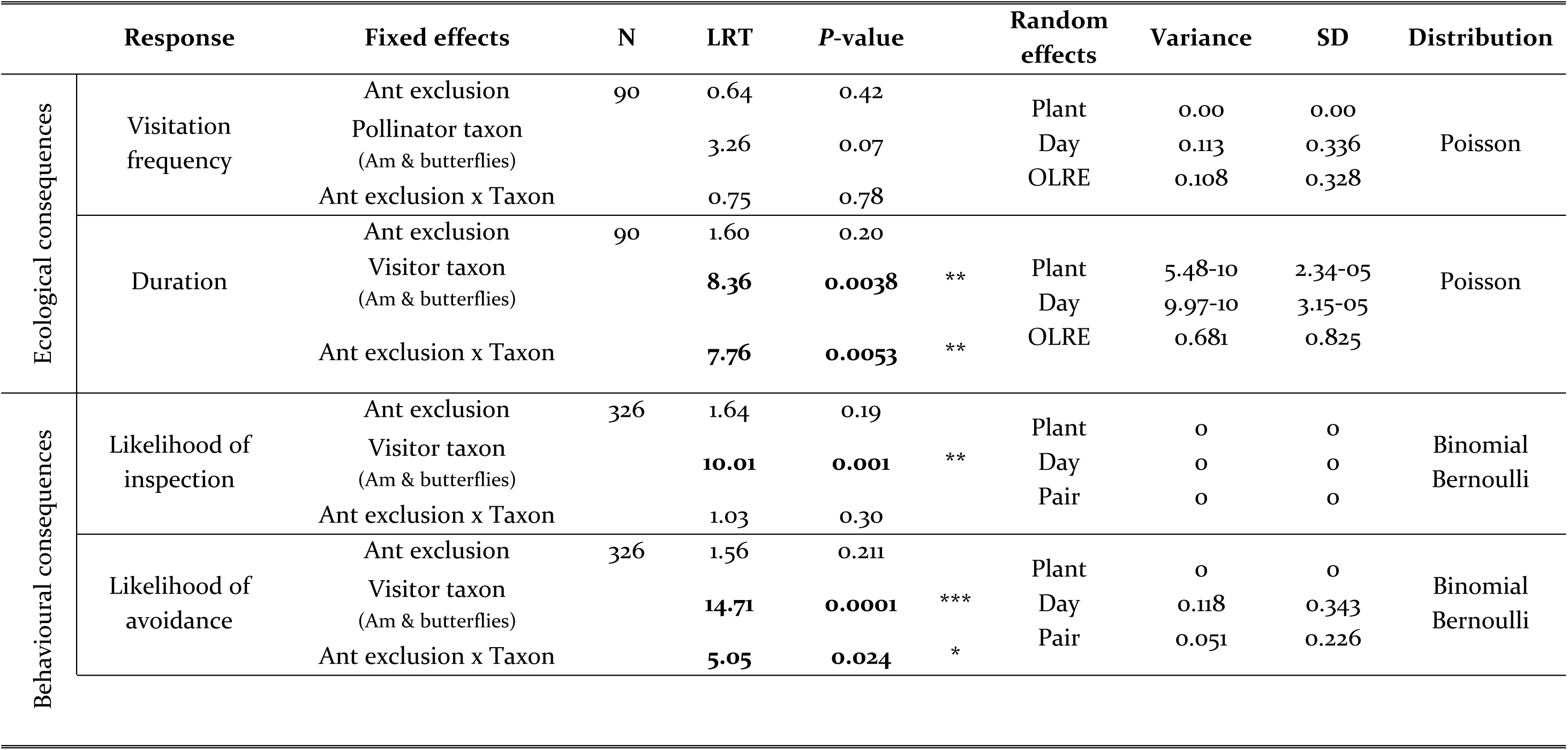
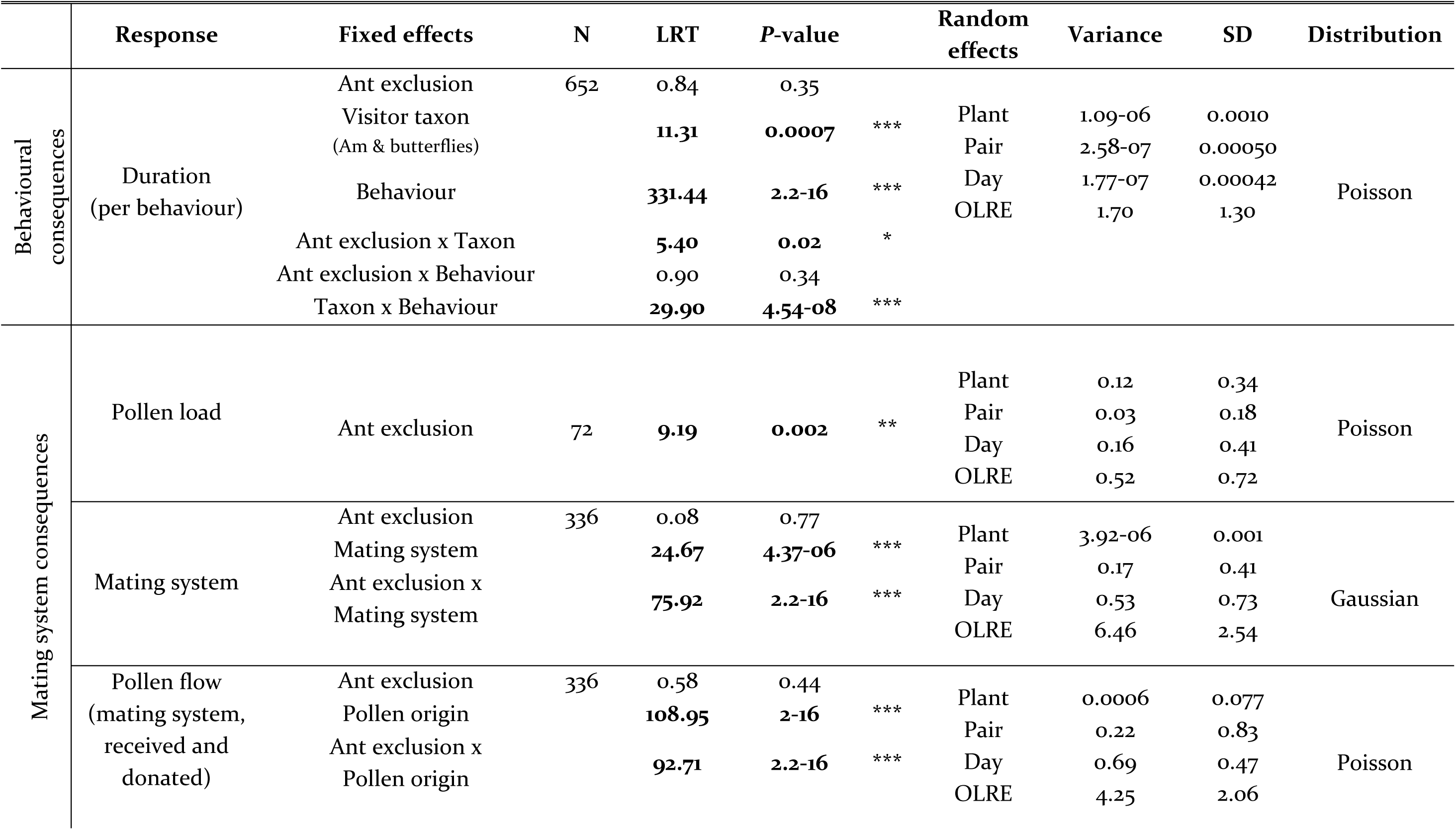
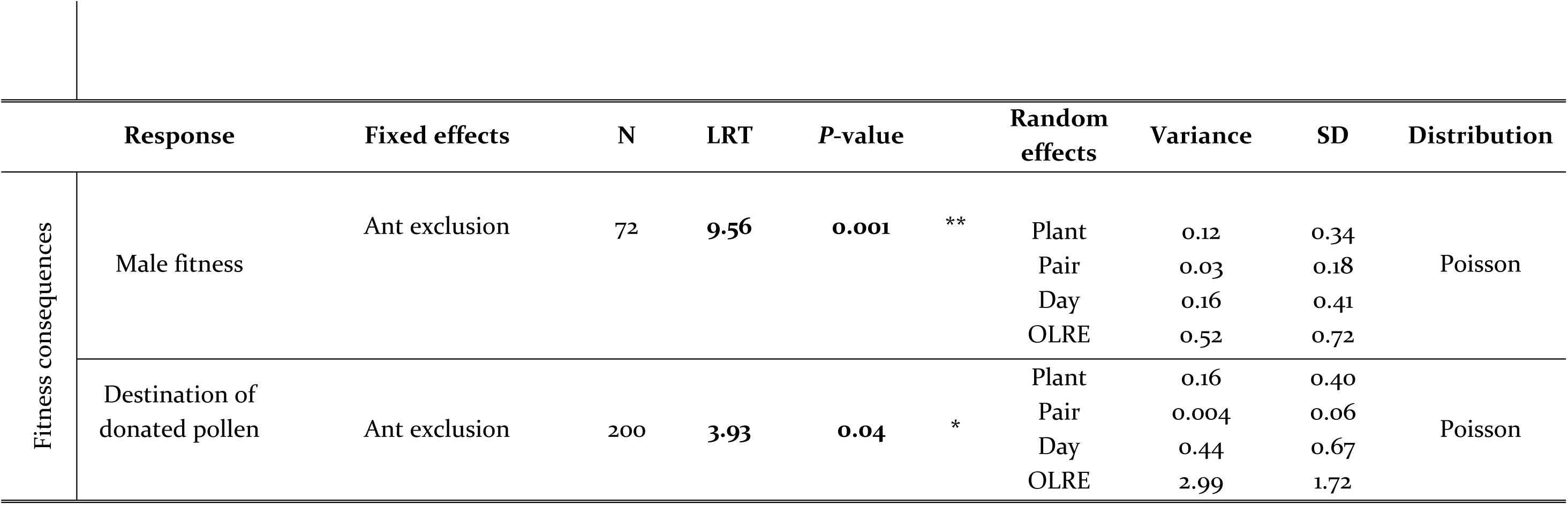
Model statistics testing the costs of ant patrolling on *Turnera velutina*’s pollination biology, including ecological, behavioural, mating system, and fitness consequences.

#### a) Pollinator community composition

Flower visitors regarded as potential pollinators (hereafter pollinators) were identified to one of five taxonomic categories: *Apis mellifera*, native bees, butterflies, flies, and wasps. To estimate the overall abundance of pollinators from each taxonomic group, we pooled together observations from control and ant-excluded plants and calculated the percentage of visitors from each group. Within each of these taxonomic groups, differences in the total number of visitors between control and ant excluded plants were assessed using a Pearson Chi-squared test. Because *Apis mellifera* and butterflies jointly accounted for 94% of all visitors (Table S1), only these taxonomic groups were included in all further analyses.

#### b) Pollinator visitation frequency and duration

Flower visits were scored each time a pollinator hovered over, landed and contacted the reproductive organs of a flower, and visit duration was recorded until the pollinator departed. We recorded visitor identity and considered revisitation events. Visitor abundance was estimated as the number of individual visitors per taxa landing on flowers of a particular plant. For instance, a visitor that landed, hovered, and landed again in another flower was registered as two visits from one visitor. Ant patrolling effects on visitation frequency were tested using a Poisson mixed model. The effect of ant patrolling on visit duration was tested using a Poisson mixed model.

#### c) Pollinator behavior

All pollinator visits were allocated to one of two behavioural categories following Villamil *et al.* (2018): inspection (defined as a pollinator approaching a flower without landing) or contact (landing on the flower). The effect of ant patrolling on the likelihood of pollinators displaying inspection behaviours was tested with a binomial mixed model, considering the presence or absence of inspection behaviours as the response variable. The effect of ant patrolling on pollinator deterrence was tested using a binomial mixed model. Pollinator deterrence is here defined as the absence of contact behaviours following an inspection behaviour. For every pollinator that displayed an inspection behaviour, we recorded the presence or absence of contact behaviours and fitted this as a binomial response variable. For instance, if a pollinator hovered over a flower, without landing inside it, we would record a zero as the response variable. The effect of ant patrolling on the duration of each type of behaviour was tested using a Poisson mixed model, splitting observations into inspection or contact behaviours. The total duration of each behaviour (inspection or contact) per visitor was fitted as the response variable.

### Plant mating system and pollen transfer dynamics

The effect of ant patrolling on pollen transfer and its consequences on plant mating system was assessed by counting differentially dyed pollen grains on focal flower stigma squash slides under a light microscope. The effect of ant patrolling on stigma pollen load, defined as the total number of pollen grains received per stigma, was tested using a Poisson mixed model (Table 1).

Pollen colour allowed us to identify pollen grains received from either the same flower (selfing), another flower within the same plant (geitonogamy), the other focal plant in the same array (outcrossing), or another un-dyed plant (outcrossing). The number of pollen grains from each origin (selfing, geitonogamy or outcrossing) was divided by the total number of pollen grains stigma (pollen load) to determine the proportion of pollen from each mating system source. Proportional data were transformed to normality using the logit transformation, with infinite numbers resulting from impossible quotients replaced by zeros. The effect of ant patrolling on the mating system was tested using a linear mixed model, fitting the proportion of pollen from each mating system as the response variable (Table 1).

Pollen transfer dynamics were analysed using five categories to describe the mating system and the pollen origin (hereafter referred to as MSPO; Fig. 1a). These categories summarise pollen grains received from and donated to every possible pollen source identifiable in this experiment as follows: (i) received/donated to the same flower (selfing), (ii) received from another flower from the same plant (geitonogamy received), (iii) received from the reciprocal focal plant (outcrossing pair received), (iv) donated to the reciprocal focal plant (outcrossing pair donated), (v) or received from another plant from the same species (outcrossing unknown received). The effect of ant patrolling on pollen flow dynamics was tested using a Poisson mixed model (Table 1), fitting as the response variable the number of pollen grains in each of the five MSPO categories.

### Male plant fitness

The number of pollen grains donated per flower was an estimate for male plant fitness and quantified as number of pollen grains from each flower donated to focal stigmas. The total number of pollen grains from satellite flowers on the same plant was divided by five, to obtain the mean number of pollen grains donated per flower. The total number of pollen grains from the other focal plant in the same array was divided by six (1 focal + 5 satellite flowers). The effect of ants on male fitness was estimated using a Poisson mixed model fitting as the response variable the number of pollen grains donated per flower in control or ant-excluded plants.

The effect of ant patrolling on the destination of the pollen grains donated per flower was tested using a Poisson mixed model, fitting number of pollen grains as the response variable. We only contrasted the number of pollen grains donated by focal or satellite flowers, as every plant had exactly six flowers because floral display was controlled for in our experimental design. Pollen donated by unknown plants was excluded as the number of donor flowers was unknown, and hence pollen grains donated per flower cannot be estimated.

## Results

### Pollinator visitation: composition, frequency and duration

We recorded 967 floral visitors, of which 853 belonged to taxa we regarded as potential pollen vectors (hereafter pollinators) because they were observed contacting male and female plant sexual organs (Table 1), although experimental analyses of their efficiencies as pollen vectors are required. Butterflies and honeybees accounted for more than 80% of all floral visitors and > 94 % of potential pollinators (Table S1). Ant exclusion did not significantly influence the community composition of pollinators visiting *T. velutina* flowers (*X*^*2*^ = 1.42, df = 4, *P* = 0.84; Fig. 2).

**Figure 2.**
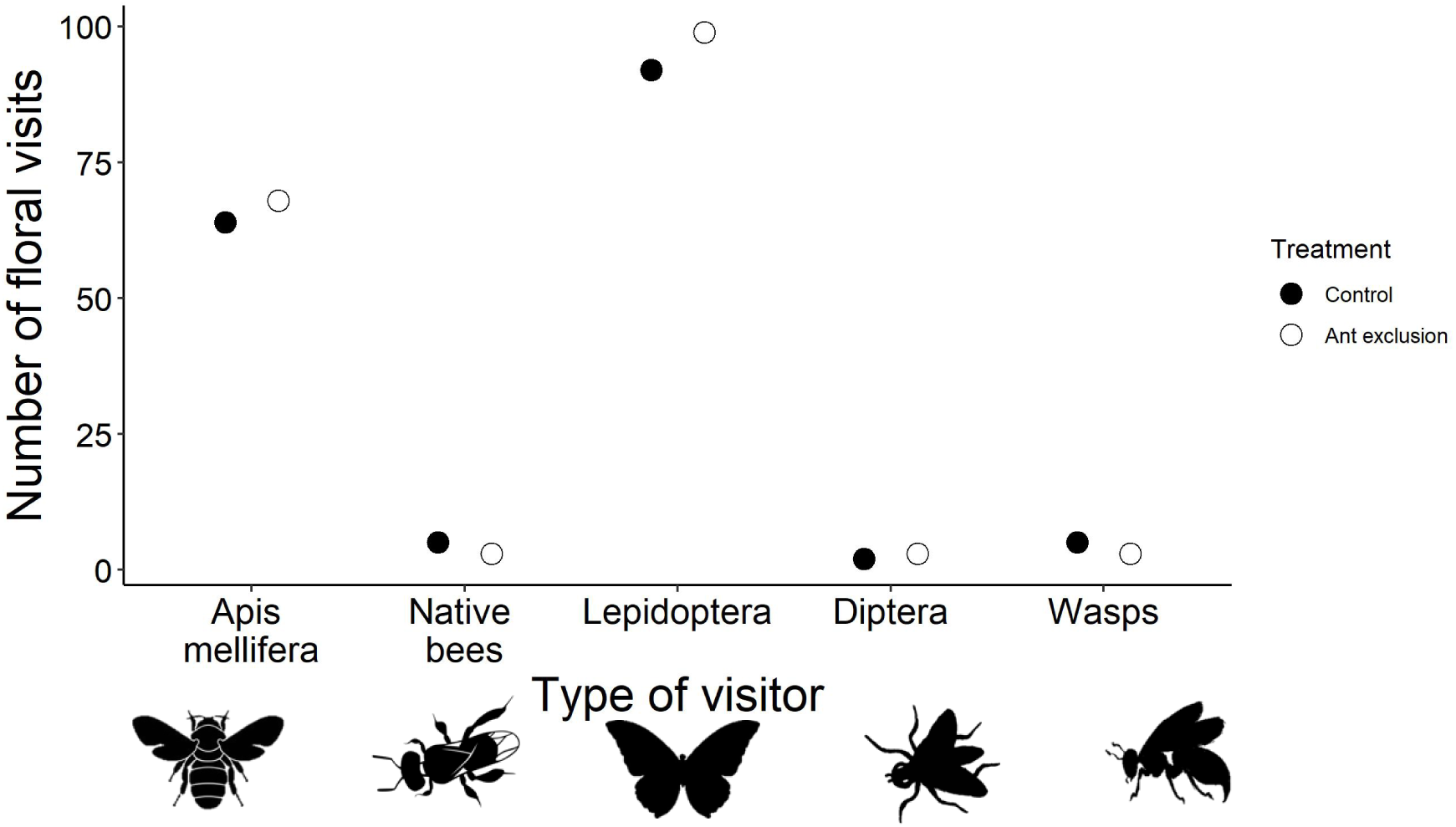
Composition of floral visitors in *Turnera velutina* plants with and without ant patrolling.

Visitation frequency did not differ significantly between pollinator types, or between control and ant-excluded plants (Fig 3a, Table 1). The effect of ant exclusion on visit duration varied across pollinator taxa, as indicated by the significant interaction term (Fig. 4b, Table 1). While flower visits by honeybees were twice as long in ant-excluded plants (*Z* = 2.45, *P* = 0.05; Table 1; Fig. 4b), there was no significant effect of ant patrolling on butterflies (*Z* = 1.07, *P* = 0.70; Table 1; Fig. 4b).

**Figure 3.**
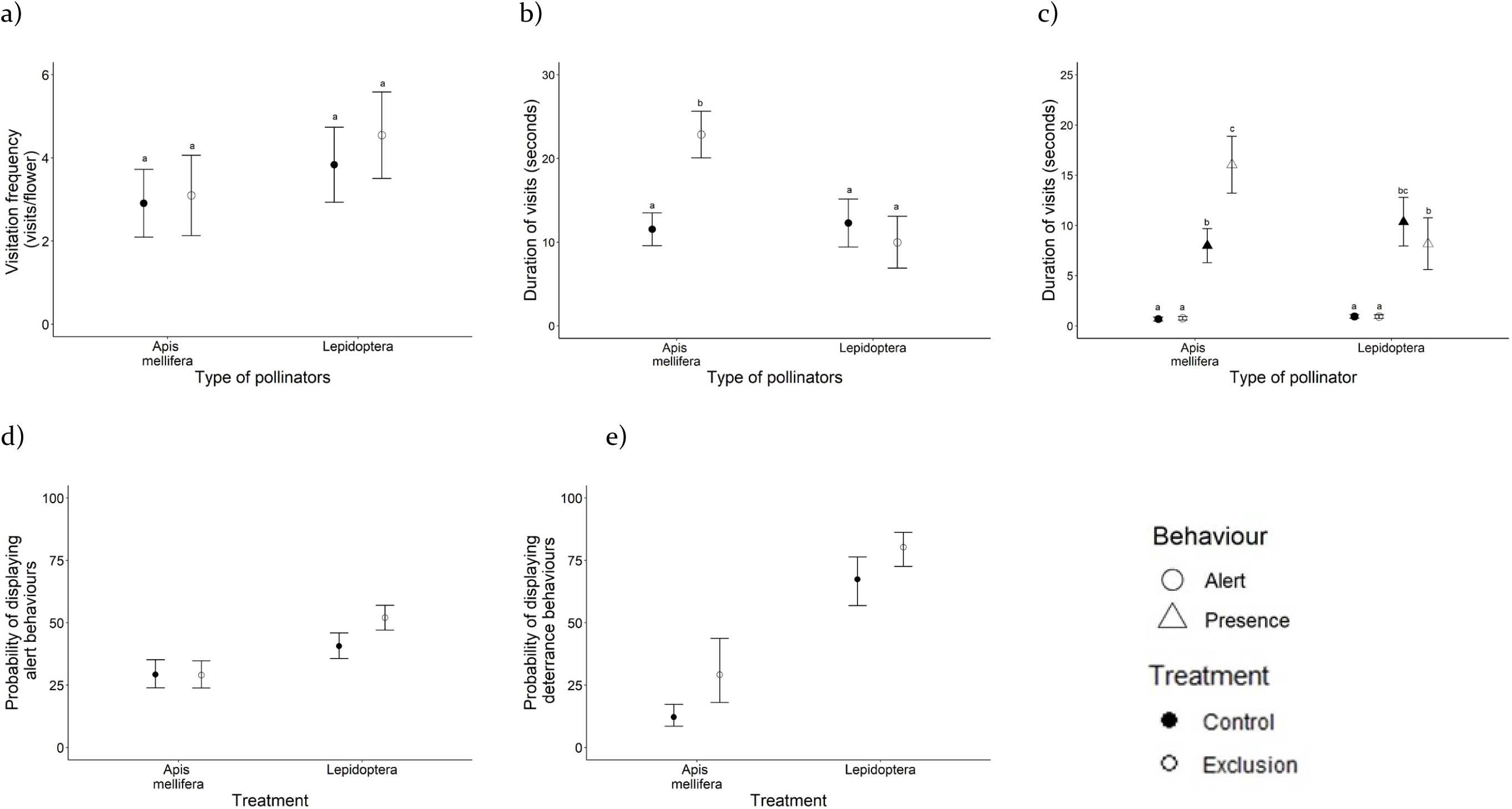
Effects of ant patrolling on pollinator visitation by *Apis mellifera* and native butterflies on *Turnera velutina* flowers on control plants with ant patrolling (black), and ant excluded plants (white). (a) Pollinator visitation frequency, (b) visit duration, and (c-e) pollinator behaviours affecting (c) the time spent displaying alert (circles) or contact/presence behaviours (triangles), (d) the display of alert behaviours, (e) the likelihood of deterrence.

**Figure 4.**
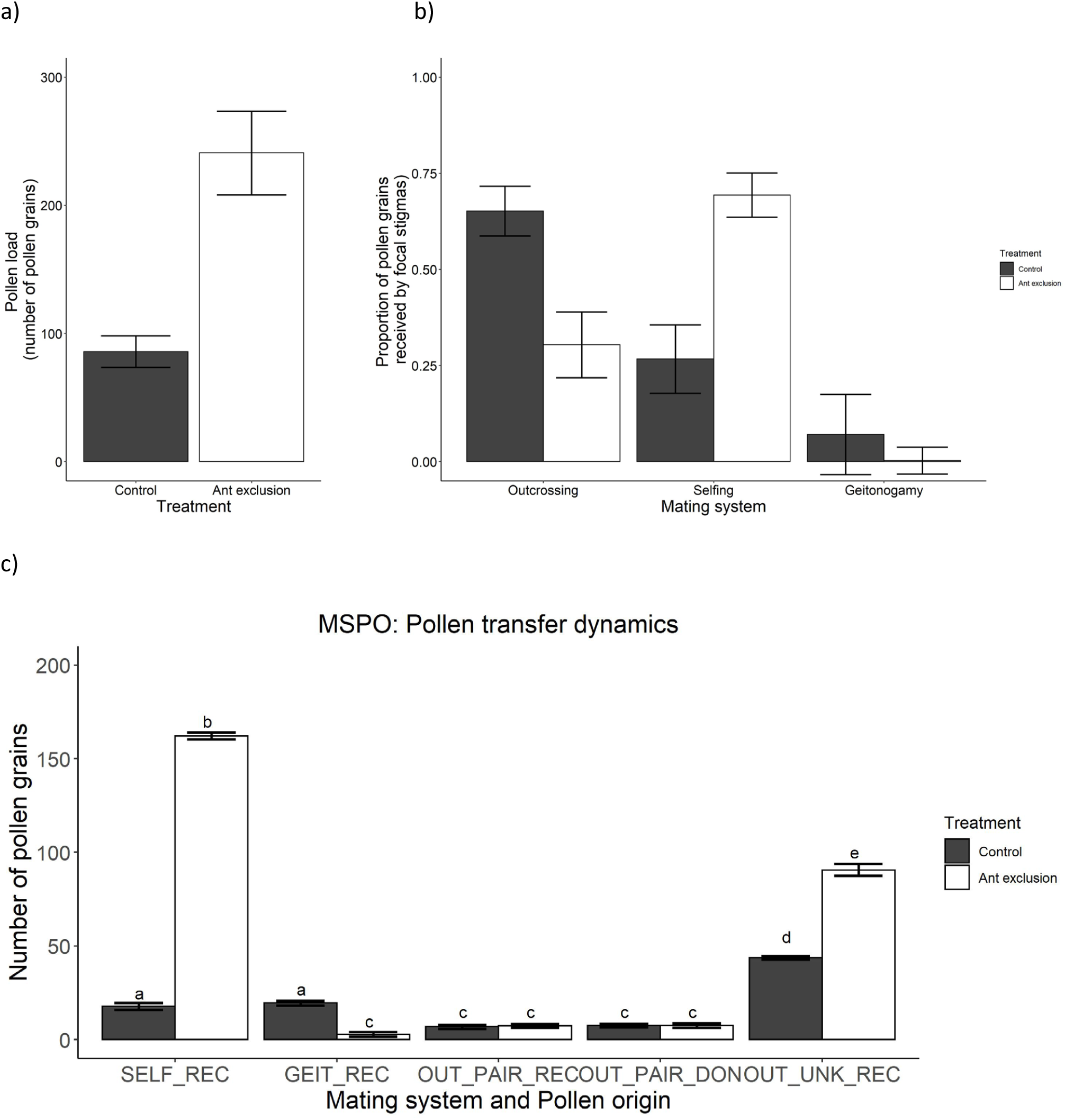
Effects of ant patrolling on the (a) pollen load, (b) mating system rates and (c) pollen flow (origin and destination) in *Turnera velutina* showing mean ± se for control (black) and antexcluded (white) plants.

### Pollinator behaviour

Inspection behaviours differed significantly between pollinator taxa (Table 1, Fig. 3c), with butterflies being on average 15% more likely to display inspection behaviours than *Apis mellifera* (Fig. 3c). However, ant exclusion did not affect this behaviour in either pollinator group (Table 1, Fig. 3c). Avoidance differed significantly between pollinator taxa, and the two taxa differed in their responses to ant guards (significant ant exclusion × pollinator taxon interaction; Table 1). Ant exclusion increased avoidance behaviour in butterflies, but decreased it for *Apis mellifera* (Fig. 3d), resulting in butterflies being deterred from landing on flowers following inspection three times more frequently than *Apis mellifera* (butterflies: 27%, *Apis mellifera*: 8.5%; Fig. 3d, Table 1).

When visit duration was split between inspection and contact behaviours, the effect of ant exclusion on visit duration differed between pollinator taxa, and behaviours (Fig. 3e, Table 1). Ant exclusion significantly increased the duration of *Apis mellifera* contact visits (*Z* = 2.96, *P* = 0.05), increasing the time bees spent inside flowers, but did not affect the time butterflies spent inside flowers (contact behaviours: *Z* = −2.34, *P* = 0.25), or the duration of inspection behaviours by either pollinator (*Apis mellifera*: *Z* = – 0.48, *P* = 0.99; butterflies: *Z* = −1.73, *P* = 0.65). Both pollinator groups spent longer periods displaying contact behaviours than inspection behaviours, regardless of the ant exclusion treatment (Table 1, Fig. 3e).

### Plant mating system and pollen transfer dynamics

Pollen load per stigma was significantly higher in ant-excluded plants (Fig. 4a, Table 1), with focal stigmas on ant-excluded plants receiving on average 155 more pollen grains than stigmas on control flowers (control: 85 ± 12; ant exclusion: 240 ± 32 (mean ± se); LRT = 9.19, *P* = 0.002; Table 2, Fig. 4a). The proportion of pollen grains from each mating system category differed significantly within and between plant treatments (Fig. 4b, Table 1). In particular, ant exclusion halved outcrossing rates, reduced geitonogamy 33-fold, and tripled selfing rates (Table 1, Fig. 4b).

Ant exclusion increased the number of selfing and allogamous pollen grains from non-focal plants received by stigmas, but reduced the number of geitonogamous pollen grains (Table 1, Fig. 4c). But overall, ant exclusion had no effect on the number of pollen grains received and donated by flowers between reciprocal pair plants (OUT_PAIR_REC: control *vs*. exclusion: Z = 0.31, P = 1.00; Fig. 4c).

### Plant fitness

Ant exclusion increased male fitness, assessed as the number of pollen grains donated per flower, from 27.2 ± 6.54 pollen grains in control plants to 163 ± 23.9 in ant-excluded plants (mean ± se) (Fig 5a). Ant exclusion, mating system, and their interaction all had significant effects on the number of pollen grains donated per flower to different destinations (Table 1). Most of the pollen donated per flower was received on the same flower’s stigma as selfing pollen, regardless of the exclusion treatment (Fig. 5b). Furthermore, ant patrolling had no significant effect on the number of pollen grains donated to the reciprocal pair plant.

**Figure 5.**
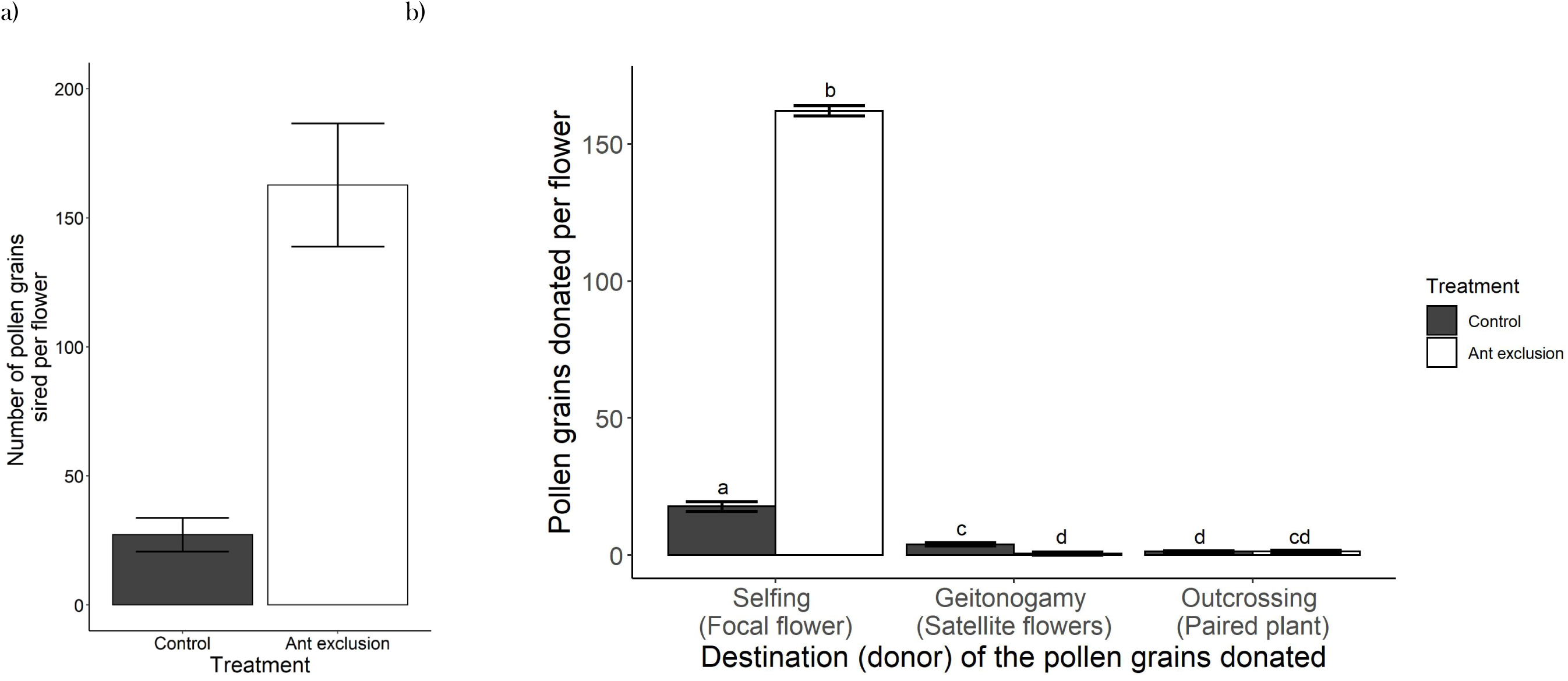
Effects of ant patrolling on male fitness, showing (a) average number pollen grains fathered per flower, and (b) the destination of the pollen grains donated per flower (mean ± se) for control (black) and ant excluded (white) plants.

## Discussion

This study provides a comprehensive picture of the interaction between myrmecophily and pollination by showing the ecological and behavioural effects of ant patrolling on pollinators and their cascading effects on plant mating system and fitness. Despite previous experimental evidence in this system suggesting direct ant-pollinator conflicts (Villamil *et al.* 2018), and contrary to our expectations, excluding ants from plants did not affect pollinator community composition (Fig. 2), visitation frequency, pollinator avoidance or inspection behaviours (Fig. 3). However, ant exclusion increased pollinator visit duration (Fig 3), pollen load, male fitness, and selfing rates (Fig. 4). To our knowledge, this is the first evidence that ant patrolling can affect the host plant mating system and male plant fitness.

### How do ants affect the plant mating system and fitness?

Ant exclusion doubled the time *Apis mellifera* spent inside flowers and increased pollen load on stigmas by 150%, but did not affect visitation frequency. These findings are consistent with the pollinator relocation hypothesis, which suggests ants can mildly deter pollinators leading to equally frequent but shorter visits that enhance pollen transfer. Furthermore, ant exclusion promoted a switch in the mating system from outcrossing to selfing (Fig. 4b). The increase in the time *Apis mellifera* spent inside flowers may underlie the increased selfing rates observed in ant-excluded plants: by foraging longer on pollen and nectar bees are likely to transfer more pollen from the anthers to the stigmas within a flower. Longer contact visits by *Apis mellifera* in the absence of ants may also be responsible for the increased male fitness, if longer visits allow *Apis mellifera* to collect and transport more pollen grains. Hence, ants cause behavioural changes in pollinator visitation dynamics that have cascading effects on the host plant mating system, and ultimately influence male and female fitness.

### Effects of ant patrolling on anti-predatory responses and efficiency

Anti-predatory responses in pollinators vary depending on the pollinator and predator taxa involved (Romero *et al.* 2011), but few studies have documented how different floral visitors respond to ant patrolling (Ness 2006; Ohm & Miller 2014; Carper *et al.* 2016) and how different ant partners affect pollinators (Ness 2006; Miller 2007; Ohm & Miller 2014; Villamil *et al.* 2018). Several hypotheses have been made regarding how ants may differentially affect each pollinator depending on body size and lifestyle (social or solitary) (Clark & Dukas 1994; Abbott & Dukas 2009) (Romero et al. 2011). Overall, predation risk by ant patrolling in *T. velutina* was not strong enough to affect pollinator composition or increase the natural avoidance and inspection rates of pollinators. Our behavioural results are consistent with the pattern revealed in a meta-analysis by Romero *et al.* (2011) showing that pollinator lifestyle (social *vs*. solitary) is not a good predictor of anti-predatory sensitivity. In our study, ant patrolling reduced visit duration in *Apis mellifera*, but not in butterflies (Fig. 3b, Table 1). Our results show avoidance behaviours differ between pollinator taxa, a pattern consistent with previous findings suggesting different pollinators differ in their anti-predatory response behaviours (Romero *et al.* 2011).

The net effect of defensive mutualists on the host plant pollination and fitness can vary depending on whether predators deter efficient pollinators or inefficient visitors (Romero & Koricheva 2011). For example, guarding ants decreased plant fitness when they attacked efficient pollinators in *Ficus pertusa* (Moraceae: Bronstein 1991) and *Opuntia imbricata* (Cactaceae:Ohm & Miller 2014), but had positive effects in *Banistriopsis malifolia* (Malpighiaceae:Alves-Silva *et al.* 2013) where wasps protected flowers from predation without deterring efficient pollinators. In *T. velutina* butterflies were more abundant than bees, but changes in the behavioural patterns of bees, and not of butterflies, (Fig. 3) seem to be driving changes in plant mating systems (Fig. 4) and fitness (Fig. 5).

### Effects of ant patrolling on plant mating system and pollen transfer

Although the deleterious effects of selfing and geitonogamy have been well described for many species (Waser & Price 1991; de Jong *et al.* 1992; Lloyd 1992), the decomposition of pollen on stigmas into donor components (intraflower or selfing, intraplant or geitonogamy, interplant or outcrossing) has rarely been performed, and its importance remains underappreciated (de Jong *et al.* 1993; Wu *et al.* 2018). We assessed the effects of ant patrolling on plant mating system decomposing pollen transfer based on its origin and fate finding that in this self-compatible species, ant exclusion shifted the plant mating system from predominantly outcrossing to predominantly selfing, reducing geitonogamy (Fig. 4b). In bisexual flowers self-pollination can be mediated by pollinators (Wu *et al.* 2018) and we suggest that this is the case for *T. velutina* where ant patrolling reduced selfing rates by affecting pollinator visitation behaviour, reducing the time bees spent inside flowers.

When contrasting our results from the mating system analyses (which show the proportion of pollen from different origins received by a stigma) with data on pollen transfer (showing counts of pollen grains from different donors) it becomes evident that the increase in pollen load in the absence of ants is driven by selfing and allogamous pollen from other undyed, surrounding, *T. velutina* plants. The number of selfing pollen grains on ant-excluded stigmas was much higher, likely driving the change in the mating system from outcrossing to selfing. Yet, ant exclusion also increased the number of allogamous pollen grains received by the stigmas. The significant increase in the number of geitonogamous pollen grains received by ant-patrolled stigmas (Fig. 4c) is consistent with the pollinator relocation hypothesis, which proposes that ant patrolling mildly deters pollinators, causing them to move to a nearby flower and hence increasing the rate of geitonogamy in the plant (Altshuler 1999; Romero & Koricheva 2011). Our results suggest that ants contribute to maintaining outcrossing in the self-compatible *T. velutina*.

### Effects of ant patrolling on plant fitness

A complete understanding of the effects mutualists and antagonists have on plant fitness requires the assessment of both female and male fitness components, because the magnitude and direction of the effects may differ between plant sexual functions (Schaeffer *et al.* 2013; Carper *et al.* 2016). For instance, male fitness is more constrained by the number of mates reached than female fitness; and pollinator behaviour affects pollen transfer, with longer visits increasing pollen export and pollen deposition on stigmas (Carper *et al.* 2016). Consequently, we expect male fitness to be more susceptible to changes in pollinator behaviour (Krupnick & Weis 1999; Schaeffer *et al.* 2013; Carper *et al.* 2016). To date, very little has been done to assess the effect of ant patrolling on female fitness, and we are not aware of any study assessing the effects of ants on male plant fitness. Our experimental design allowed us to quantify the consequences of ant patrolling on male reproductive fitness (pollen grains donated per flower) and infer potential effects on female fitness (progeny quality). Ant exclusion resulted in a six-fold increase in the number of pollen grains donated per flower (Fig. 5a), suggesting that guarding ants may hinder male fitness.

Ant exclusion changed the pollen destination, given that most pollen was donated towards selfing, which contrasts with flowers from ant-patrolled flowers which donated a quarter as much pollen to themselves (Fig. 5b). Hence, ant patrolling in *T. velutina*’s may increase female fitness by promoting outbred seeds and so increasing the offspring quality. Our findings contrast with previous studies showing negative effects of ant patrolling on female plant fitness. In *Opuntia imbricata* ant patrolling decreased seed count by 30% and seed mass by 16% (Ohm & Miller 2014); and in *Heteropterys physophora* ants consumed floral buds, deterred pollinators, reduced pollen transfer and fruit set in buds that escaped ant predation (Malé *et al.* 2012). Furthermore, our findings exemplify how non-pollinators insects interacting with plants and their pollinators may have contrasting effects on female and male fitness components, highlighting the importance of considering both sexual functions.

This study provides a comprehensive picture of the interaction between myrmecophily and pollination by showing the ecological and behavioural effects of ant patrolling on pollinators and its cascading effects on plant mating systems that lead to fitness consequences. Contrary to our initial prediction, ant patrolling benefited plant fitness by reducing pollinator visit duration, which promoted pollinator relocation, and led to a reduction in selfing and an increase in outcrossing rates. Although ant patrolling reduced pollen load and male fitness, far from having an ecological cost on the host, ant patrolling seems to be another mechanism – along with herkogamy and pollinator attracting features (Sosenski *et al.* 2016)– to promote outcrossing in this self-compatible species with hermaphroditic flowers. This study contributes towards our understanding on how non-pollinators can shape plant mating systems. We provide the first evidence of the role of patrolling ants on determining plant mating systems and male plant fitness by taking a multispecies approach on plant-animal interactions.

## Acknowledgements

We thank Aura Larsson and Alberto Herrera for their help during fieldwork, Rubén Pérez-Ishiwara for logistical assistance, Jarrod Hadfield for valuable inputs towards the statistical analyses, and Ally Phillimore for comments which greatly enhanced previous versions of the manuscript.

